# RepairSwitch: simultaneous functional assessment of homologous recombination vs end joining DNA repair pathways in living cells

**DOI:** 10.1101/2023.01.17.524235

**Authors:** Ruchama C. Steinberg, Jianyong Liu, Ajay M. Vaghasia, Hugh Giovinazzo, Minh-Tam Pham, Dimitri Tselenchuk, Roshan Chikarmane, Michael C. Haffner, William G. Nelson, Srinivasan Yegnasubramanian

## Abstract

DNA repair pathways are frequently defective in human cancers. DNA double strand breaks (DSBs) are most often repaired by either homologous recombination (HR) or non-homologous end joining (NHEJ). Alterations in repair pathways can indicate sensitivity to therapeutic agents such as PARP inhibitors, cisplatin, and immunotherapy. Thus, functional assays to measure rates of HR and NHEJ are of significant interest. Several methods have been developed to measure rates of HR or NHEJ; however, there is a need for functional cell-based assays that can measure rates by both major DNA DSB pathways simultaneously. Here, we describe the RepairSwitch assay, a flow cytometry assay to assess rates of HR and NHEJ mediated repair of Cas9 programmed DSB simultaneously using a novel fluorescence switching reporter system. The assay exhibits low background signal and is capable of detecting rare repair events in the 1 in 10,000 range. We demonstrate the utility of RepairSwitch by measuring the potency of inhibitors of ATM (KU-60019, KU-55933), DNA-PK (NU7441), and PARP (Olaparib) on modulating DSB repair rates in HEK293FT cells. The selective ATM inhibitor KU-60019 inhibited HR rates with IC50 of 915 nM. Interestingly, KU-60019 exposure led to a dose responsive increase in rates of NHEJ. In contrast, the less selective ATM inhibitor KU-55933, which also has activity on DNA-PK, showed inhibition of both HR and NHEJ. The selective DNA-PK inhibitor NU7441 inhibited NHEJ efficiency with an IC50 of 299 nM, and showed a dose responsive increase in HR. The PARP inhibitor Olaparib showed lower potency in modulating HR and NHEJ. We next used the RepairSwitch assay to assess how pharmacological and genetic inhibition of DNA methyltransferases (DNMT) impacted rates of HR and NHEJ. The DNMT inhibitor decitabine reduced HR, but increased rates of NHEJ, both in a dose responsive manner, in both HEK293FT and HCT116 cells (IC50 for HR of 187 nM and 1.4 uM respectively). Knockout of DNMT1 and DNMT3B increased NHEJ, while knockout of DNMT3B, but not DNMT1, reduced HR. These results illustrate the utility of RepairSwitch as a functional assay for measuring changes in rates of DSB repair induced by pharmacological or genetic perturbation. Furthermore, the findings illustrate the potential for one DNA repair mechanism to compensate in part for loss of another. Finally, we showed that inhibition of DNMT can lead to reduction of HR and increase in NHEJ, providing some additional insight into recently observed synergy of DNMT inhibitors with PARP inhibitors for cancer treatment.

## INTRODUCTION

There are many types of DNA damage including bulky adducts, base mismatches, insertion, deletions, single strand breaks, and double strand breaks, among others. The repair of DNA damage is essential to maintain genome integrity and prevent cell death; as such, each type of DNA damage requires one of several unique repair pathways to resolve the damage effectively. These pathways, comprised of specialized proteins necessary for each distinct avenue of repair, often harbor mutations associated with specific tumor types. Many cancers result from abnormalities in DNA repair pathway proteins, such as mutations in BRCA1/2, associated with breast, ovarian, and prostate cancers among others, and those in mismatch repair genes, which are associated with several cancers including colorectal cancer.

Double strand breaks, which are particularly genotoxic, can be repaired by multiple mechanisms, including homologous recombination, non-homologous end joining, single strand annealing, and microhomology end joining. However, the two mechanisms utilized most often to repair DNA double stranded breaks (DSBs) are homologous recombination (HR) and non-homologous end joining and other end joining pathways (NHEJ). These pathways differ from one another in a number of ways. HR is a conservative process that uses a homologous template, namely the sister chromatid in S or G2 phases of the cell cycle, or the homologous chromosome, i.e. the alternate allele when a sister chromatid is unavailable, to repair the damaged section of DNA. End joining pathways, including NHEJ, in contrast, are error prone and often mutagenic. NHEJ involves ligation of ends of the broken DNA, often adding or deleting nucleotides prior to ligation, which results in deletions and frameshifts. Therefore, further insight into the dynamics of these pathways is necessary to obtain an improved understanding of the underlying mechanisms that regulate repair.

In addition, many cancers harbor defects in the HR or NHEJ machineries. Defects in HR are largely driven by mutation or epigenetic silencing of genes like BRCA1 and BRCA2, as well as ATM, ATR, and RAD51. Familial forms of breast, ovarian, and pancreatic cancers have also been associated with loss of function mutations in these proteins, and these proteins are also frequently mutated in prostate cancers. There have been efforts to catalogue many of the mutations found in these genes; however, for many, we still don’t know what functional impact they have on repair. Synthetic lethality approaches have also been explored as therapeutic approaches for HR-deficient tumors. One important success from such synthetic lethality strategies is the use of drugs that inhibit poly-ADP ribose polymerase (PARP) enzymes in cancers with HR defects, particularly BRCA1/2 mutations. Cancer causing mutations in NHEJ seem to be less common, although patients have been identified with mutations in DNA-PK, Ligase4, and XRCC4 (Sishc and Davis 2017). Additionally, downregulation of Ku70/80 and DNA-PK has been found in several cancers, which could be one mechanism of promoting chromosomal instability and driving tumor progression. However, there is still significant uncertainty about whether a given mutation may be functional in impacting DNA repair pathways, or whether cells may have functional loss of repair even in the absence of currently known mutations, for example as a result of epigenetic alterations, or drug exposures.

As such, the impact of HR and NHEJ pathway perturbation on rates of repair is an active area of study. Several methods have been developed to address this question; however, most require the use of multiple tools to fully assess rates of repair by each pathway individually. For example, the widely used fluorescence-based assay, utilizing the DR-GRP reporter, can be used to assay the frequency of HR at a chromosomal DSB (Jasin 1996; Andrew J. Pierce and Jasin 2005). It does this through the use of a nonfunctional GFP vector, interrupted by a I-SceI locus, that when cut and repaired by HR will yield a functional GFP protein. However, it cannot determine the frequency of NHEJ or any other type of repair without the use of PCR-based assays in conjunction (Vriend, Jasin, and Krawczyk 2014). Additional constructs utilizing I-SceI have been developed to assess the frequency of NHEJ as well as HR; however, as both constructs result in the reconstitution of GFP expression, they cannot be used to determine rates of repair by each pathway within a single sample (Seluanov, Mao, and Gorbunova 2010).

The Traffic Light assay is one such system, developed to gain a better understanding of repair pathway choice at DNA breaks and to evaluate the efficiency and outcome of nuclease induced genome editing (Certo et al. 2011). This system is designed to produce a DSB at the I-SceI nuclease cleavage site within a nonfunctional GFP that can then be repaired by NHEJ or by HR with the addition of a repair template that is lentivirally transduced. In this system, if HR is used to repair the locus a functional EGFP protein will be produced. However, if NHEJ is used to repair the locus, and results in a +3 frameshift, the construct will produce a functional mCherry protein instead. This is achieved by way of a T2A self-cleaving peptide that enables the mCherry to be cleaved from the nonfunctional GFP and escape degradation. One limitation of this system is that +3 frameshifts represent a third or less of all mutagenic events that can occur during repair by NHEJ. The efficiency and outcome of the repair by HR vs NHEJ varies with increased lentiviral nuclease delivery. Lastly, one more caveat of this system is that it requires the generation of reporter cell lines and cannot be easily extended to new cell systems in a modular fashion.

Therefore, to address the unmet need for an efficient system that overcomes many of these caveats, here we have developed a novel and modular cell-based assay to assess rates of HR and NHEJ simultaneously using flow cytometry of living cells that we call RepairSwitch. The assay takes advantage of CRISPR/CAS induced DNA double strand breaks directed at an eGFP template, followed by HR mediated switching to blue fluorescence protein, or error-prone end-joining mediated mutation and loss of eGFP, all in the background of cells constitutively expressing the red fluorescence protein dsRED. The constitutive expression of the dsRED protein allows gating for cells that have definitely incorporated the reporters, and have maintained enough viability to express this protein even after introduction of the DSB, allowing for differentiation between loss of eGFP due to mutation induced by end joining, vs loss of eGFP due to loss of cell viability. The application of switching from eGFP to BFP has been used previously to assess efficiency of CRISPR/CAS genome editing (Glaser, McColl, and Vadolas 2016), but has not been used to assess rates of DNA repair in cells. Using our novel RepairSwitch assay, we assess the impact of ATM and DNA-PK inhibition on rates of repair, and show that both HR and NHEJ can act in a compensatory manner upon inhibition of proteins in either pathway. Further, by inhibiting proteins whose role in repair is not restricted to either pathway, we establish the assay’s utility in determining repair pathway selectivity. An improved understanding of these DNA repair machinery players may lead us to novel therapeutics and new avenues of cancer prevention.

## MATERIALS AND METHODS

### Plasmid Construction

Vectors for this reporter assay were constructed using a lentiviral construct containing CMV-DsRed and UBC-EGFP on a pHAGE backbone purchased from Addgene (plasmid #24526). This lentiviral construct then underwent site directed mutagenesis using the Agilent QuikChange II kit, introducing an AgeI cut site at the c-terminus of the UBC promoter (3897-3901) via 3 base changes. The plasmid was then restriction digested by AgeI and BamHI to remove the UBC promoter via gel extraction. DNA encoding the EF1a promoter was PCR amplified using NEB Phusion HF DNA Polymerase Kit, attaching AgeI and BamHI restriction sites to the 3’ and 5’ ends, respectively, and was subsequently digested. This amplicon was then cloned into the pHAGE backbone using cohesive ligation via a Quick Ligase Kit.

Plasmids expressing DsRed, EGFP, and BFP individually were constructed as compensation control for flow cytometry using this backbone. The DsRed only plasmid was constructed via restriction digestion by BamHI and ClaI and subsequent gel extraction of the reporter assay construct, to remove the EF1a-UBC region. A short, ∼30bp oligo, purchased from IDT with BamHI and ClaI restriction sites on the 5’ and 3’ ends, respectively, was then digested and used to cohesively ligate the ends via Quick Ligase Kit. To create the EGFP only construct, the reporter assay construct was restriction digested using SpeI and BamHI followed by gel extraction to remove CMV-DsRed. The resulting backbone was treated with Mung Bean Nuclease and a Quick Ligase Kit was used to circularize the plasmid via blunt end ligation. This plasmid then underwent site directed mutagenesis using the Agilent QuikChange II Kit to create the BFP only plasmid (within EGFP: C197G_T199C).

### Viral Packaging

Lentiviral packaging vectors pMDLg/pRRE (5ug), pRSV-Rev (2.5ug), and pMD2.G (2.5ug) were used to package viruses of all assay vectors (10ug of vector). FuGENE HD was used to transfect packaging vectors and assay constructs into HEK293FT cells for viral production in a ratio of 3:1 FuGENE to DNA. Media was replaced 4-16 hours post transfection. Virus was collected 48-hours post transfection and then 12-hours later, combining both batches. Virus was then spun down, aliquoted, and stored at −80 C.

### Transduction, MOI, and FACS

Spinfection protocol was used to transduce cell lines, adding 8ug/ml polybrene to the growth medium along with virus followed by centrifugal spin for 2 hours before being placed in the incubator. Media was replaced 24-hours post transduction and flow cytometry and FACS was performed after 72-96 hours. MOI of 0.3 was assessed using viral dose curve and flow cytometry. FACS was used to sort positive cell populations.

### Gene Targeting

A Cas9/gRNA vector targeting EGFP was constructed using the LentiCRISPRV2 backbone and their established protocol for gRNA production. The EGFP-targeting gRNA sequence was designed using ChopChop.

### HR Repair Template

ssDNA oligo containing BFP homologous template to GFP was purchased from IDT and resuspended to a 1ug/ul concentration. This repair template is ∼100bp in length, protected at 5’ and 3’ ends with two consecutive phosphorothioate bonds. Methylated versions of this template were also purchased containing methyl groups at all available CpGs, seven sites in total. Methylated versions of the aforementioned ssDNA oligo template were also purchased from IDT and resuspended to a 1ug/ul concentration. These methylated oligos contain methyl groups at all available CpGs, seven sites in total. This repair template is also ∼100bp in length and protected at 5’ and 3’ ends with two consecutive phosphorothioate bonds.

### Cell Culture

HEK293FT cells were obtained from ThermoFisher Scientific (R70007) and subsequently cultured in DMEM, a high glucose medium supplemented with 10% FBS. HCT116 DNMT1 KO and DNMT3B KO cell lines were gifts from the Baylin Lab and were cultured in McCoy’s growth medium supplemented with 10% FBS. CWR22RV1 cells and CWR22RV1 ATM −/− cells were obtained from Clovis and cultured in RPMI-1640 growth medium supplemented with 10% FBS.

### Cell Line Transfections with CRISPR/Cas9 Reagents and HR Repair Template

All cell lines were plated in 12-well plates at 2.5E5 cells/well in their respective growth mediums. The growth medium was then replaced after 24-48 hours and the cells were transfected with 1ug of CRISPR/Cas9 gRNA construct, 1ug of HR repair template, or both gRNA and HR repair template (2ug total) in combination using FuGENE HD (Promega) at a FuGENE to DNA ratio of 3:1. The DNA was mixed with Opti-MEM (50ul Opti-MEM/ug DNA) and FuGENE was added for an incubation period of 15 minutes before being added dropwise to the well. If drug is being utilized, cells are dosed immediately prior to transfection. Fresh growth medium was replaced every 48-72 hours. Cells were kept in culture for 1 week before assessment of fluorescence via flow cytometry.

### Pharmacological Inhibitors

All compounds used can be found listed in Table 1, and all were purchased from Selleck. All drugs were dissolved in DMSO as per Selleck instructions. For cell-based work drugs were diluted to a final concentration of 0.001% DMSO. All drugs were given in a dose range of 1nm to 10uM, unless toxicity was shown in Incucyte proliferation assays (S1, S2).

**Table 1.**
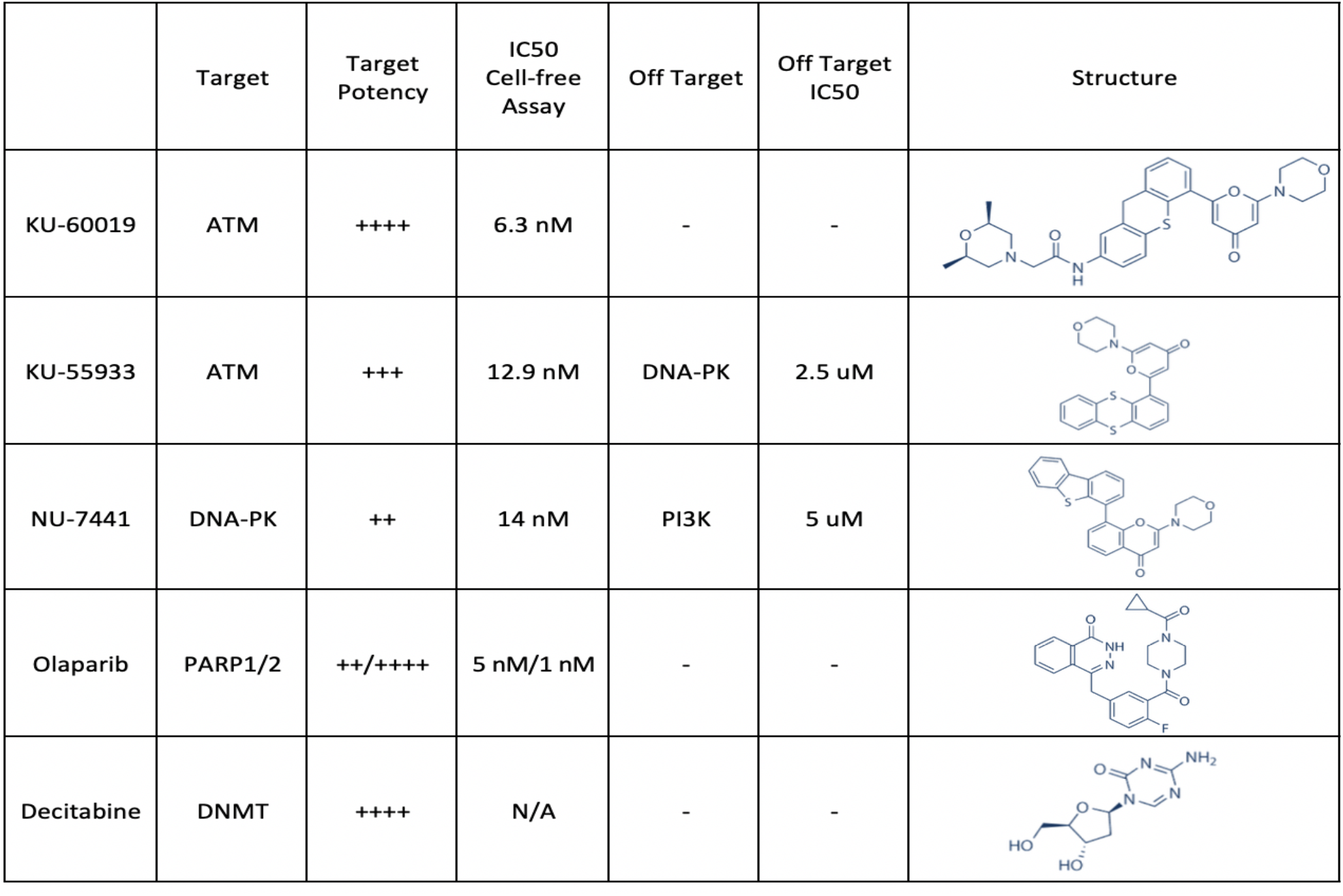
Inhibitors used in this study.

|  | Target | Target Potency | IC50 Cell-free Assay | Off Target | Off Target IC50 | Structure |
| --- | --- | --- | --- | --- | --- | --- |
| KU-60019 | ATM | ++++ | 6.3 nM | - | - |  |
| KU-55933 | ATM | +++ | 12.9 nM | DNA-PK | 2.5 uM |  |
| NU-7441 | DNA-PK | ++ | 14 nM | PI3K | 5 uM |  |
| Olaparib | PARP1/2 | ++/++++ | 5 nM/1 nM | - | - |  |
| Decitabine | DNMT | ++++ | N/A | - | - |  |

### Flow Cytometry Analysis

Medium was removed from 12-well plates and cells were detached from plate using TrypLE Express. FBS containing medium was used to neutralize TrypLE Express and cells were collected and spun down at 1000 RPM for 5 min. Cell pellets were then washed with PBS twice before resuspension in 500ul PBS and put on ice. Prior to flow of each sample, cells were resuspended and put through a cell-strainer cap on the flow tube. Cells were gated to ensure single cell population as well as high DsRed expression. For each sample, 10,000-30,000 high DsRed expressing cells were counted depending on sample concentration, and assessed for both EGFP and BFP expression. FlowJo was utilized to analyze all flow cytometry results and to produce statistics on each population.

### Visualization and Statistical Analysis

Graphpad Prism was utilized to visualize statistical results from FlowJo. Additionally, it was utilized to provide statistical analyses of p value using unpaired t-tests.

## RESULTS

### Overview of RepairSwitch assay principles

The RepairSwitch assay was developed to measure the balance of HR- and NHEJ-mediated double-strand break (DSB) repair. To accomplish this, it was necessary to design assay vectors that could be lenti-virally transduced into cells and would thereby be incorporated into the genome. We determined that a fluorescent dual reporter system, using CMV-DsRed and EF1a-EGFP, would be most effective (Fig. 1A). This design gave a strong expression of both fluorescent markers. Additionally, it allowed DsRed to serve as our transduction control to ensure vector integration at an MOI of 0.3, while EGFP served as our target for CRISPR/Cas9 induced DSB. Utilizing the CRISPR/Cas9 system as our means of DNA damage induction allowed us to ensure targeted DSB events to the fluorophore of the EGFP locus at a rate of 1 DSB per cell. Additionally, in conjunction with the administration of the CRISPR/Cas9 EGFP gRNA, exogenous single stranded DNA oligos were administered in excess and act as the repair template. In this system, the resulting fluorescence, post CRISPR/Cas9 cutting at the target locus, is dependent on the method of repair. Upon utilization of the repair template in repair via homologous recombination (HR) the resulting locus will be converted from EGFP to BFP. This is possible due to the sequence similarity of these fluorescent proteins, which differ in only two bases, resulting in the substitution of two adjacent amino acids. However, if the repair template is not utilized and repair occurs via the non-homologous end-joining pathway (NHEJ), there are two possible outcomes. Either, the EGFP fluorescence will be extinguished due to the error prone repair, or it is possible for the EGFP protein to be repaired correctly or with a silent mutation via NHEJ, thereby producing a viable fluorescent signal. However, those rare events will be indistinguishable from the DsRed+GFP+ population comprised of cells that did not experience cutting by CRISPR/Cas9. This can be due to lack of transfection by the vector or due to inaccessibility of the EGFP locus resulting from its integration in a heterochromatic region of the genome. Our schematic of the RepairSwitch assay readout (Fig. 1B) provides a concise overview of the possible fluorescent outcomes and their respective implications.

**Figure 1:**
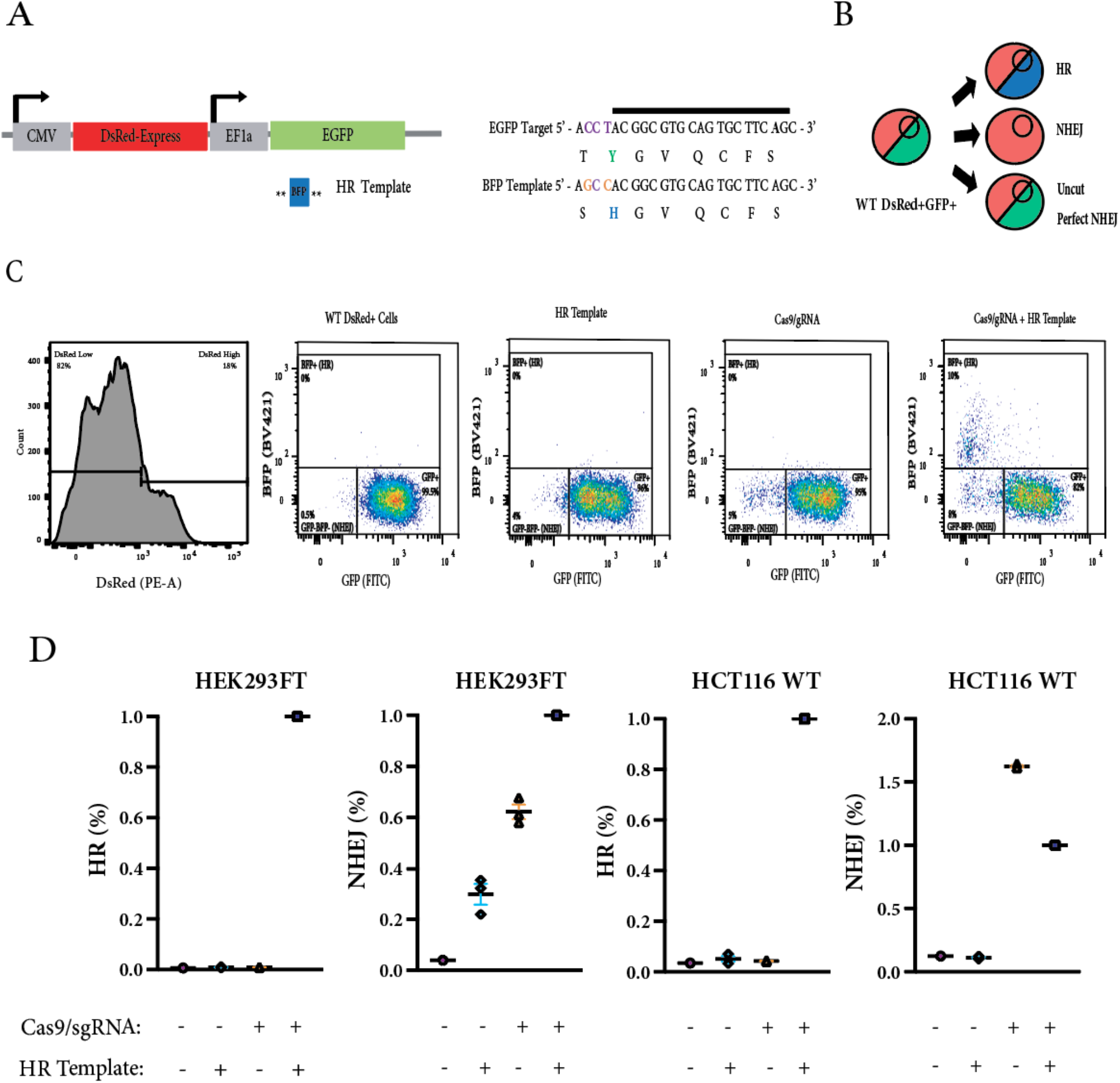
Cell based RepairSwitch assay to measure the balance of HR and NHEJ mediated DSB repair. (A) RepairSwitch assay components include: lentiviral CMV-DsRed-EF1a-EGFP vector which is transduced into cell line of interest and sorted using FACS; a LentiCRISPRv2 construct designed to target EGFP which co-expresses the Cas9 protein and the EGFP-targeting sgRNA to induce a DSB (PAM sequence in purple); and a BFP homologous recombination template protected at both ends by two consecutive phosphorothioate bonds (represented by asterisks). Sequence alignment of EGFP and BFP show that two base substitutions (orange) correspond to the two changes in amino acid sequence responsible for the shift in fluorescence from EGFP to BFP. (B) Rates of HR and NHEJ are determined by assessment of DsRed+ cells for EGFP and BFP fluorescence using flow cytometry. (C) Cells were gated on High DsRed+ expression. For each sample, 10,000-30,000 High DsRed+ cells were counted and statistical analysis was done in FlowJo and GraphPad. Representative Flow cytometry plots depict assay results and gating strategy. (D) This assay was performed in HEK239FT and HCT116 cells. In each experiment, the HR template and gRNA were each administered alone and served as assay controls.

### Assay performance with control/reference samples

This assay was performed in both HEK293FT and HCT116 cells. Flow cytometry was used to quantify results, utilizing a gating strategy designed to reduce noise while maintaining a high degree of sensitivity, detecting rare events in the 1 in 10,000 range (Fig. 1C). The results show that the expression of BFP, indicating HR has occurred, will only take place in the presence of both the CRISPR/Cas9 EGFP gRNA and the repair template (Fig. 1D). However, this is not the case for NHEJ. In HEK293FT but not HCT116 cells, EGFP signal loss is increased upon transfection of the ssDNA oligo repair template alone, indicating NHEJ has occurred at the EGFP locus without the presence of the CRISPR/Cas9 EGFP gRNA. As expected, in both cell types, the CRISPR/Cas9 EGFP gRNA alone induces loss of EGFP signal as a result of repair by NHEJ, as no repair template is available. Interestingly, the addition of both the repair template and the CRISPR/Cas9 gRNA has an additive effect on the rate NHEJ in HEK293FT cells, as seen by the higher rates of NHEJ when in combination with either of the components on their own. However, this was not observed in HCT116 cells. In HCT116 cells, the addition of both repair template and CRISPR/Cas9 gRNA results in a lower rate of NHEJ than with the gRNA alone. These results show that the RepairSwitch assay is both modular and sensitive.

### Impact of genetic modulation of repair factors on rates of repair

In order to verify the RepairSwitch assay’s efficacy in measuring rates of HR and NHEJ, we tested the impact of genetic modulation of ATM, an important and well-studied HR repair protein, in CWR22Rv1 cell lines purchased from Clovis. As expected, there was a significant reduction in the rate of HR in the ATM-/- line as compared to WT (Fig. 2A). However, surprisingly, there was also a significant increase in the rate of NHEJ in the ATM-/- cells as compared to WT (Fig. 2B).

**Figure 2:**
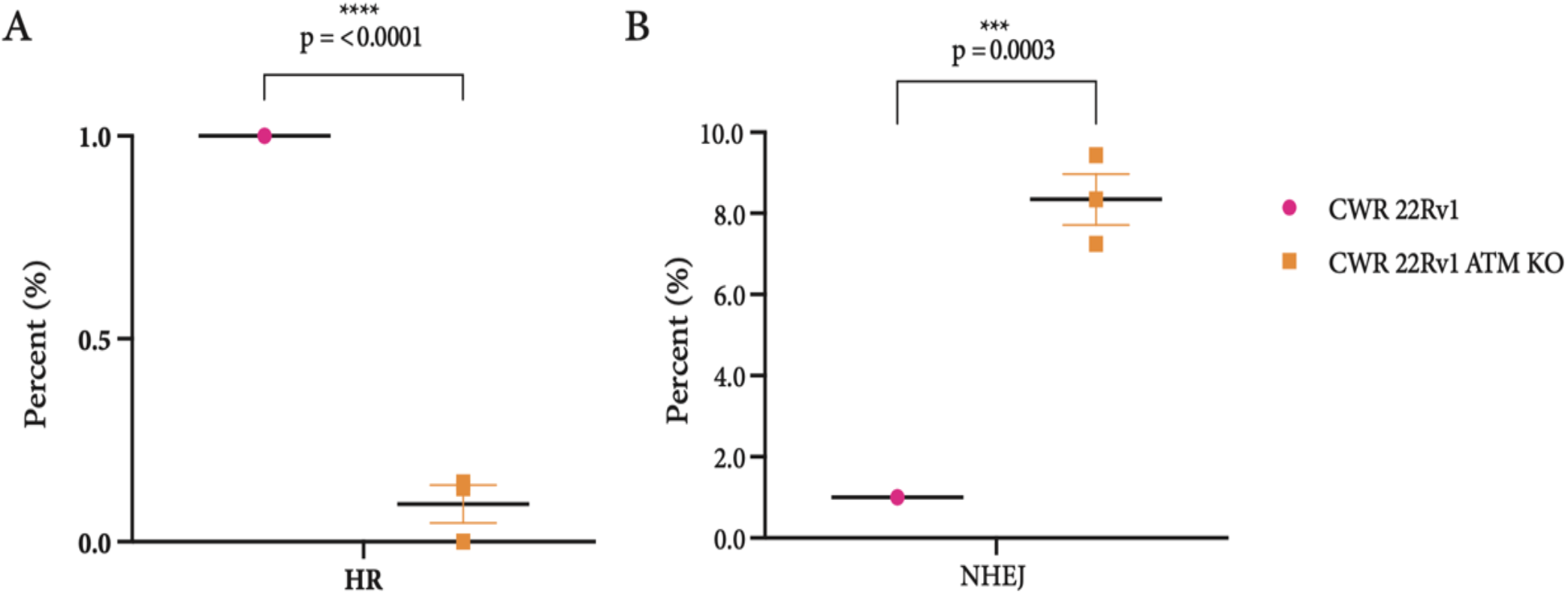
Impact of genetic modulation of repair factors on rates of repair. CWR 22Rv1 WT and CWR 22Rv1 ATM KO cells obtained from Clovis were transduced with the assay vectors and sorted prior to assay performance. Rates of HR (A) and NHEJ (B) were compared in these cell lines and statistical analysis was performed using an unpaired t-test.

### Impact of pharmacological inhibition of DNA repair enzymes on rates of repair

Utilizing a number of drugs targeting several repair factors as detailed in Table 1, we tested the impact of pharmacological inhibition on rates of HR and NHEJ in HEK293FT cells. Continuing our focus on ATM, we used two drugs that target the ATM protein, with varying degrees of potency and specificity, to ascertain whether pharmacological inhibition produced similar results to genetic manipulation of the ATM protein. KU-60019 is a potent and specific ATM inhibitor, while KU-55933 is both less potent and less specific. To ensure these drugs were non-toxic to the cells, proliferation was assayed using the Incucyte system, which showed that these drugs minimally attenuate cell growth even at higher doses. These results can be found in the supplemental material (S1. A-B). For both KU-60019 and KU-55933, HR was reduced in a dose responsive manner with an IC50 of 915 nM and 2.6 uM respectively (Fig. 3A-C). However, rates of NHEJ for the two drugs differed due to their difference in target specificity. The high degree of specificity of KU-60019 for ATM and the HR pathway, can be observed by the lack of negative impact on the rate of NHEJ with its usage. Interestingly, the rate of NHEJ can be seen to increase in a dose responsive manner as a result of KU-60019 usage. These results are consistent with our previous findings regarding the impact of genetic manipulation of ATM on the rates of HR and NHEJ. In contrast, KU-55933 has been shown to have off target effects on DNA-PK at high doses, an important component of the NHEJ pathway (Hickson et al. 2004). This is reflected in the dose responsive reduction in rates of NHEJ in following treatment with this drug (Fig. 3D).

**Figure 3:**
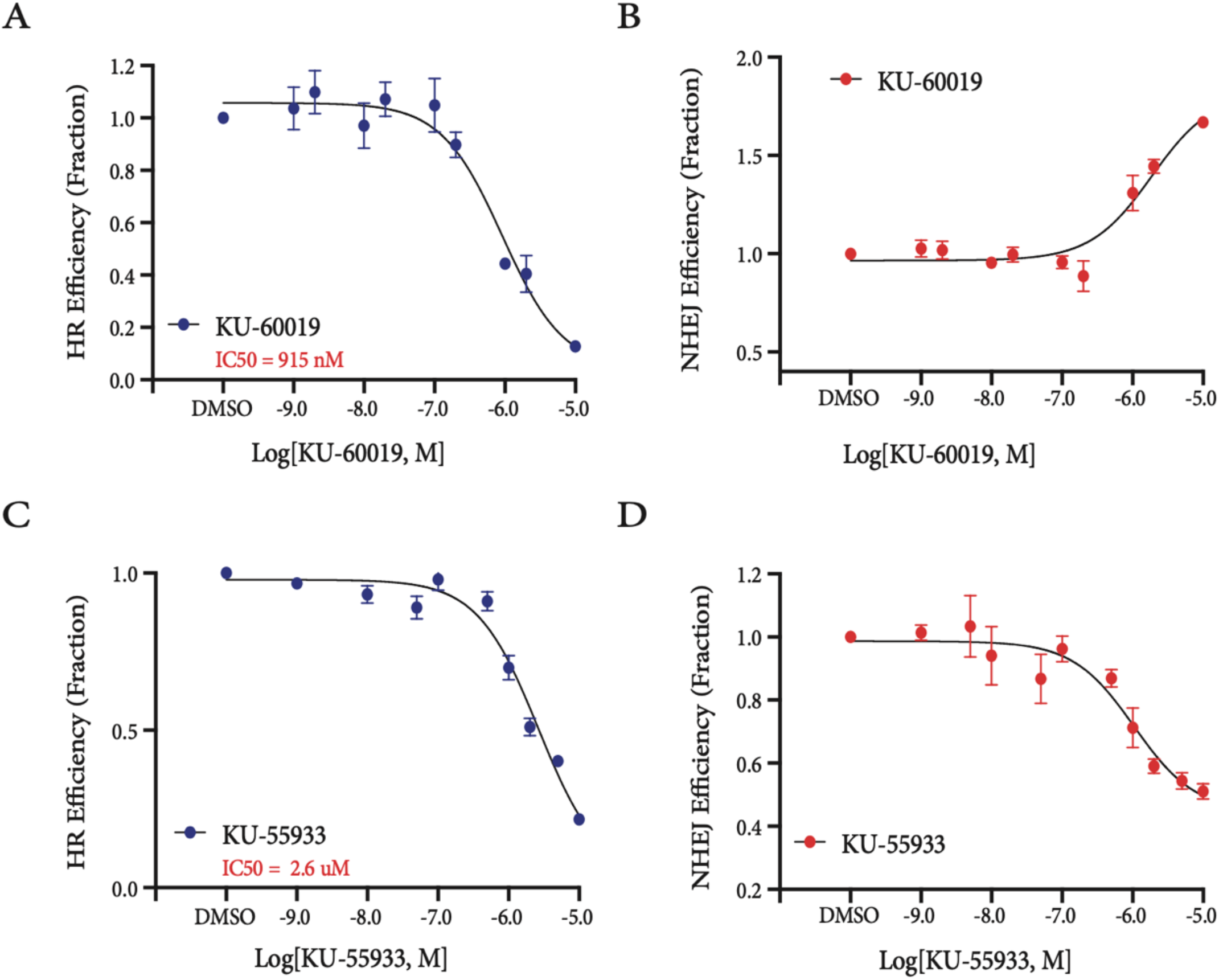
Impact of ATM inhibition on rates of repair. Rates of HR (A) and NHEJ (B) in response to inhibition of ATM using a potent and selective inhibitor, KU-60019, with an IC50 of 915nM. Rates of HR (C) and NHEJ (D) in response to inhibition of ATM using a less potent and less selective inhibitor with off targeting of DNA-PK at higher doses, KU-55933, with an IC50 of 2.6uM.

Next, we looked at the pharmacological inhibition of NHEJ components, namely DNA-PK, to assess its impact on rates of HR and NHEJ in HEK293FT cells. To do so, we utilized NU-7441, a potent and specific inhibitor of DNA-PK. Incucyte cell proliferation assays were performed to assess the toxicity of this drug, and it was found to inhibit cell growth at higher doses (S1. C). Therefore, doses that were high enough to impact cell growth were removed from this dose curve so as not to impact the data. As expected, the usage of this DNA-PK inhibitor results in a dose-dependent reduction in rates of NHEJ (Fig. 4A). However, interestingly, rates of HR also increase in a dose responsive manner. This is similar to the effect on NHEJ seen when components of HR are inhibited.

**Figure 4:**
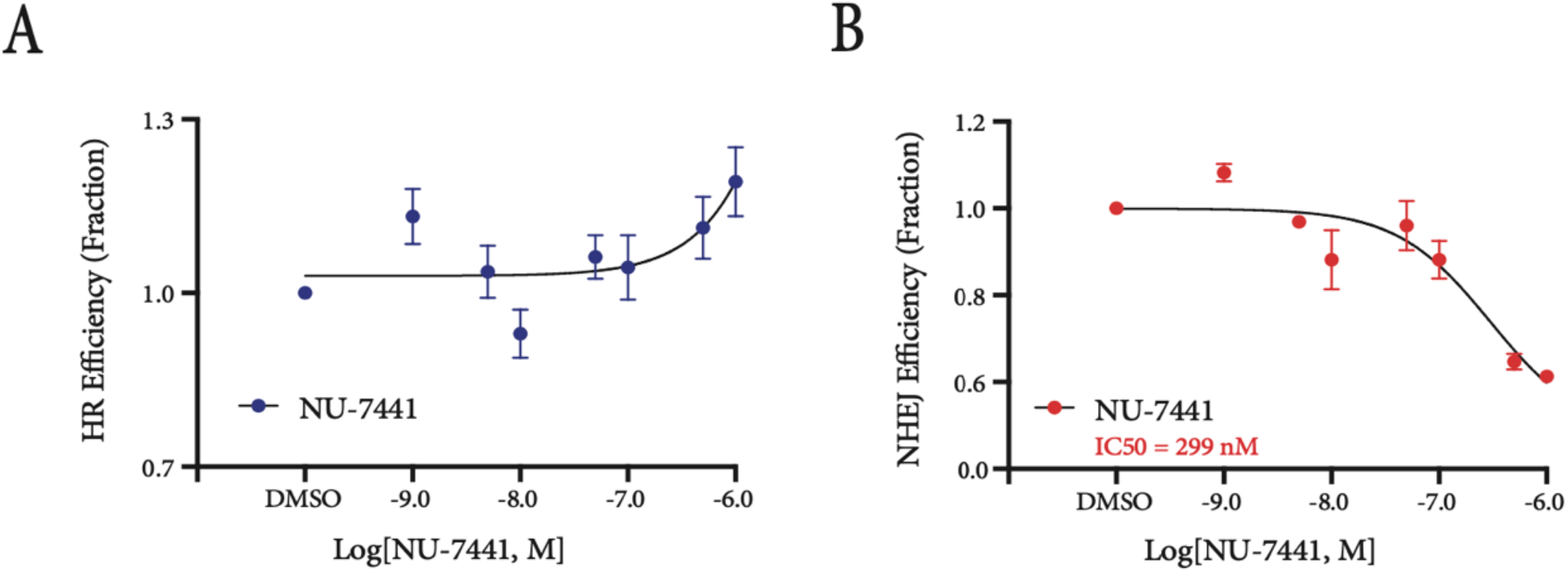
Impact of DNA-PK inhibition on rates of repair. Rates of HR (A) and NHEJ (B) in response to inhibition of DNA-PK with a potent and selective inhibitor, NU-7441, with an IC50 of 299nM.

Lastly, we used the RepairSwitch assay to provide insight into the repair function of PARP by evaluating the impact of its inhibition on rates of HR and NHEJ. These results show a subtle reduction in rates of HR and a corresponding subtle increase in the rates of NHEJ (Fig. 5A-B). Proliferation was assessed during Olaparib treatment as well, and shows it to minimally attenuate cell growth at higher doses (S1. D).

**Figure 5:**
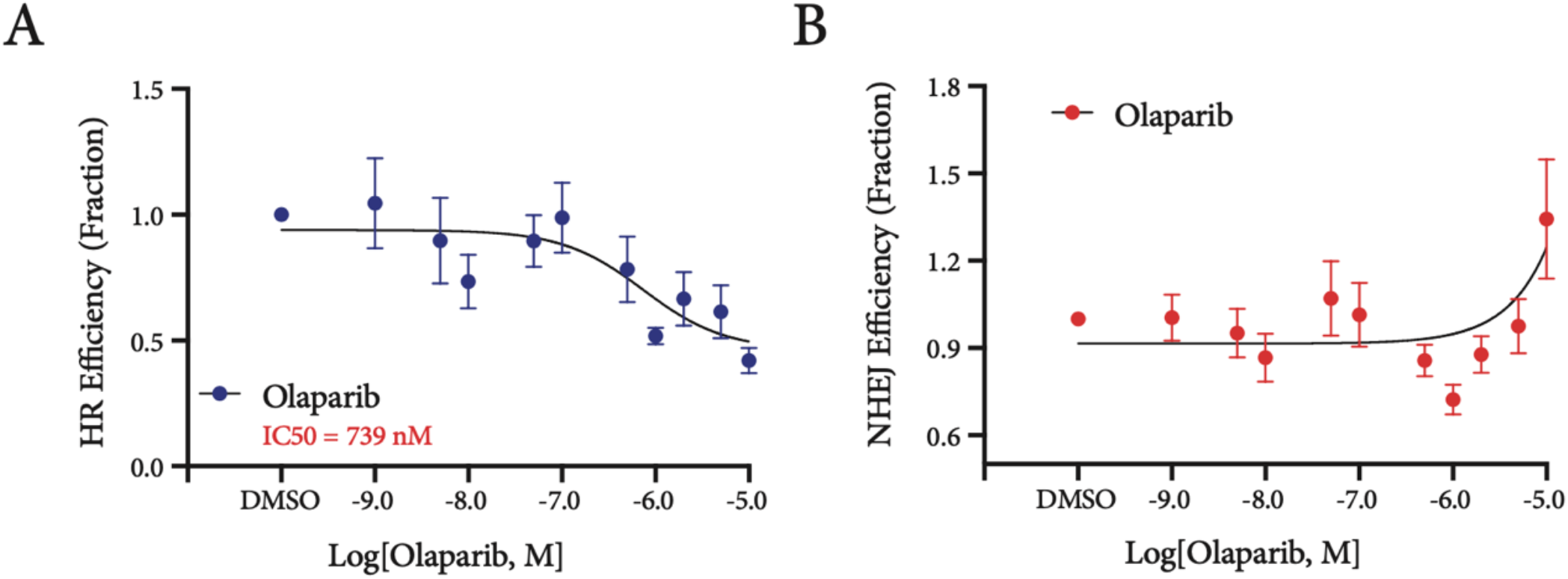
Impact of PARP inhibition on rates of repair. Rates of HR (A) and NHEJ (B) in response to inhibition of PARP with a potent and selective inhibitor, Olaparib, with an IC50 of 739 nM.

### Impact of donor template DNA methylation on rates of repair

As shown previously, the RepairSwitch Assay is designed to be versatile, and can be used to explore numerous variables that may impact rates of repair by HR and NHEJ. Here we investigated the impact of donor template DNA methylation on rates of repair. We accomplished this by using donor template DNA that is either devoid of methylation or fully methylated at all seven CpG sites within the approximately 100 bp ssOligo. This experiment was performed in both HEK293FT cells (Fig. 6A-B) and HCT116 cells (Fig. 6C-D). These results show a significant decrease in the rates of both HR (p=<0.0001) and NHEJ (p=0.0004) in HEK293FT cells when a methylated template is provided versus an unmethylated one. However, the same cannot be said of HCT116 cells, in which there was no significant difference in HR with the methylated template as compared to unmethylated. Additionally, NHEJ (p=0.0232) repair shows a small but significant increase in rates when using the methylated template as compared to unmethylated. To ensure that these results are not due to differences in how the cells recognize methylated versus unmethylated donor template DNA, statistical analysis was performed on samples that received the control template alone (Fig. 6A-D). These results showed that there were no significant differences in the rates of either HR or NHEJ upon use of either template.

**Figure 6:**
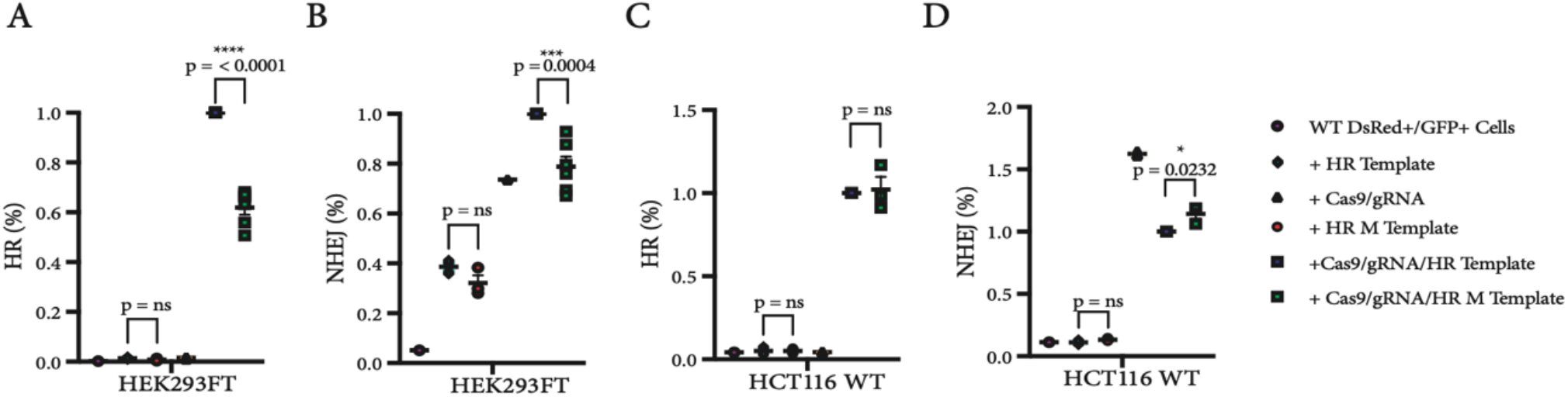
Impact of donor template DNA methylation context on rates of repair. Rates of HR and NHEJ are shown in HEK293FT cells (A,B) and HCT116 WT (C,D) cells, respectively, with both methylated and unmethylated templates along with all other assay components. Statistical analysis was performed using unpaired t-tests.

### Impact of DNMT inhibitors on rates of repair

In order to investigate the potential role of the DNA methylation machinery in repair, we used pharmacological inhibition of DNA-methyltransferases to assess the impact of loss of these pathway components on rates of HR and NHEJ. This experiment was performed in both HEK293FT and HCT116 cells. As DNA-methyltransferase inhibitors can be toxic to cells, Incucyte cell proliferation assays were performed to ensure doses at which cell growth was inhibited were excluded from our analysis (S2). Decitabine, a cytidine analog that traps DNMTs on the DNA, was used in these experiments. Our results show that treatment with Decitabine results in a dose responsive decrease in the rates of HR, and a compensatory increase in the rates of NHEJ in HEK239FT cells (Fig. 7A-B). These results were also consistent in HCT116 cells (Fig. 7C-D).

**Figure 7:**
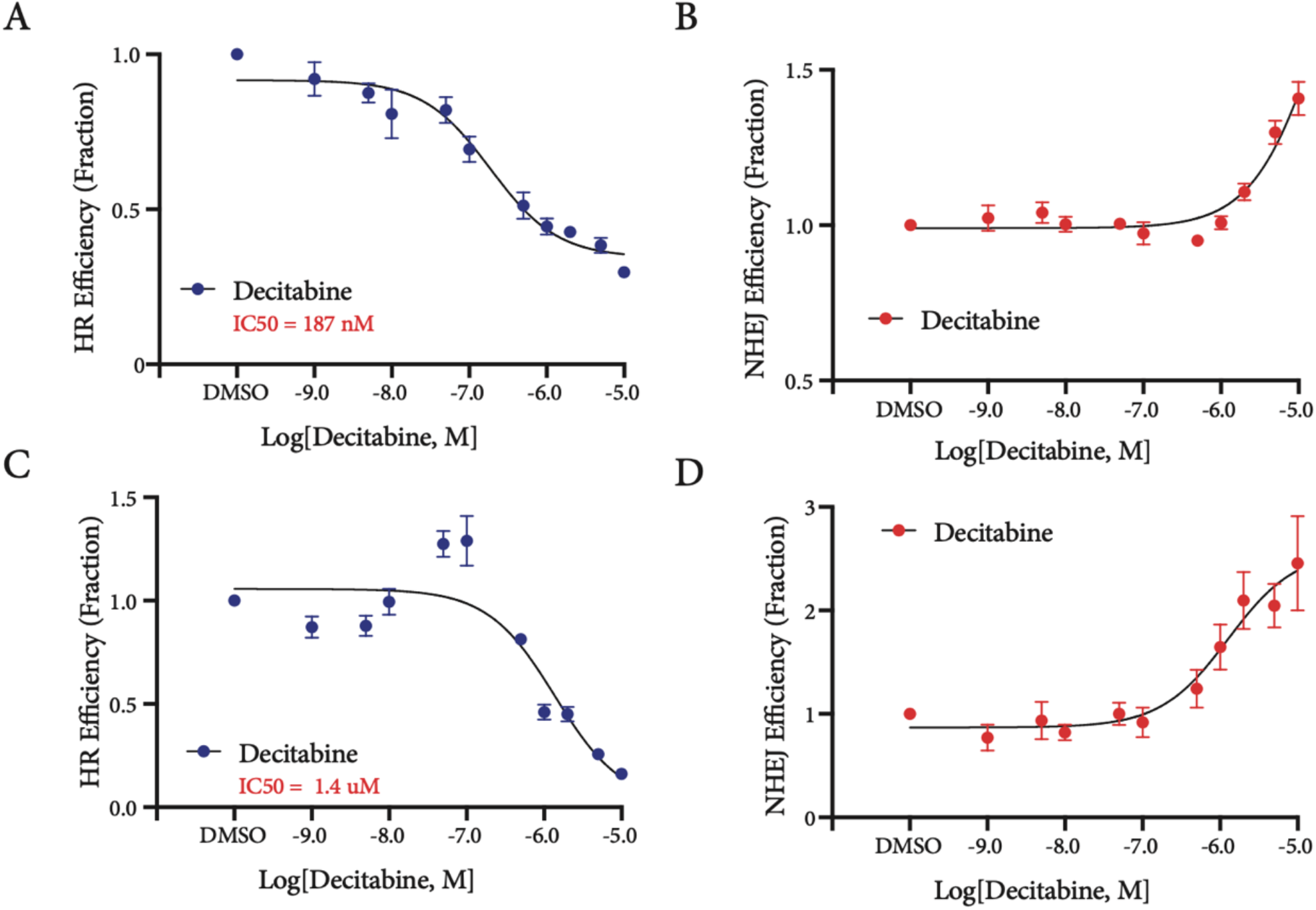
Impact of Decitabine on rates of repair. Rates of HR and NHEJ in response to Decitabine treatment in both HEK293FT (A,B) and HCT116 cells (C,D).

### Impact of genetic manipulation of DNMT genes on rates of repair

In order to get a clearer understanding of the role of DNMTs in repair, we decided to see whether these results are consistent when genetic manipulation of these components are applied. Pharmacological inhibition and genetic manipulation are complementary approaches with distinct differences. Where pharmacological inhibition of DNMTs will trap these proteins and deplete them through proteolytic degradation, genetic manipulation will remove the proteins entirely and prevent any engagement with DNA to begin with. Using HCT116 WT, DNMT1 KO, and DNMT3B KO cells, we compared rates of HR and NHEJ (Fig 8, A-B). DNMT1 KO cells had slightly increased rates of HR as compared to WT, whereas DNMT3B KO cells exhibited a drastic decrease in rates of HR as compared to WT HCT116 cells. NHEJ was significantly increased in both HCT116 DNMT KO and DNMT3B KO cells as compared to WT. This is consistent with what we see with pharmacological inhibition using Decitabine.

**Figure 8:**
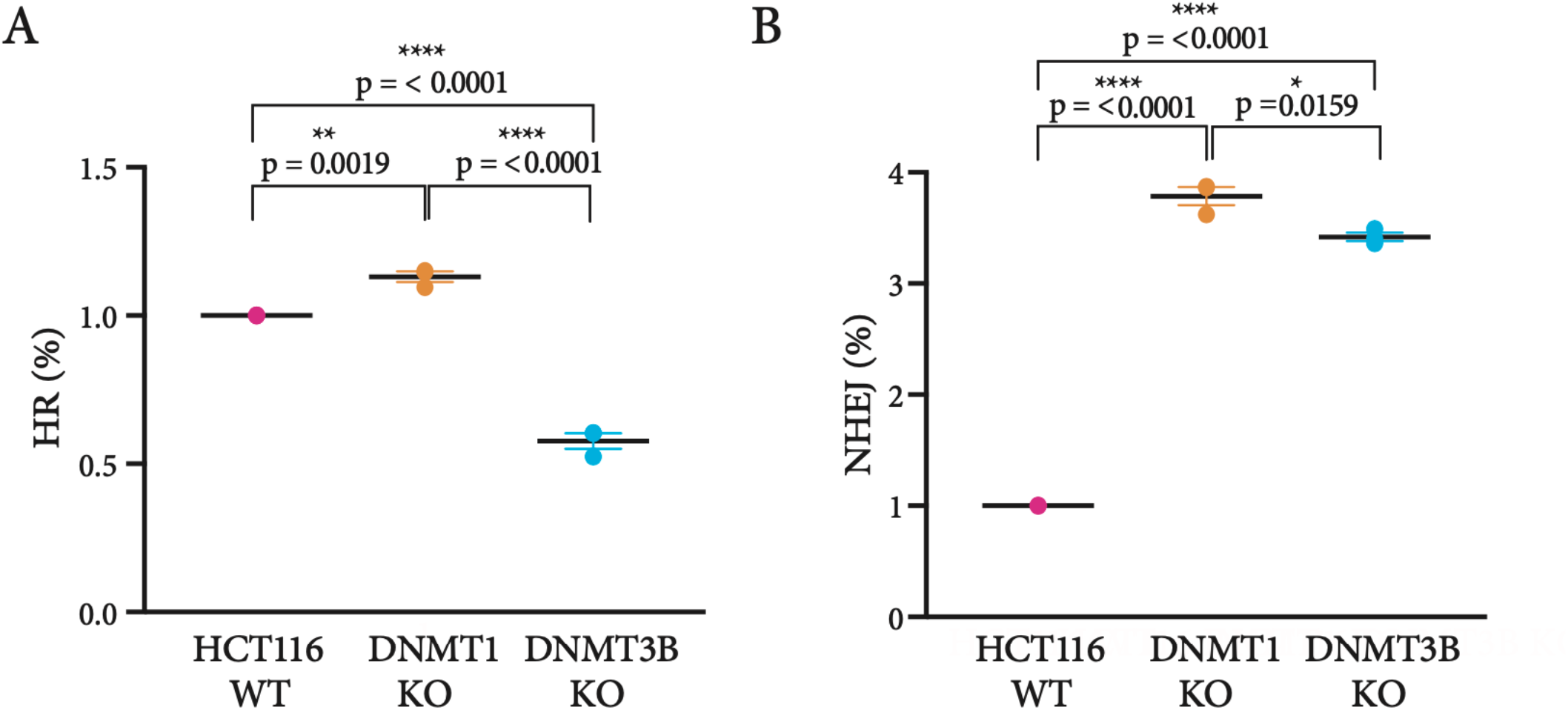
Impact of genetic manipulation of DNA methylation pathway on rates of repair. This assay was performed in HCT116 WT, HCT116 DNMT1 KO, and HCT116 DNMT3B KO cell lines and rates of HR (A) and NHEJ (B) are shown for all assay components. Rates of HR and NHEJ were compared between these cell lines and statistical analysis was performed using unpaired t-tests.

## DISCUSSION

Here we have introduced the RepairSwitch assay and explored the dynamics of DNA double strand break repair by HR and NHEJ. The RepairSwitch assay has key advantages over other assays currently in use. One such advantage being that it enables the user to obtain a rapid signal readout, using flow cytometry, of both HR and NHEJ rates within a single sample, without any loss of cell visualization. Another being that this assay utilizes the CRISPR/Cas9 system, which presents a number of advantages over the much used I-SceI nuclease, namely that it is modular and a number of gRNA’s can be used at once. Additionally, the use of CRISPR/Cas9 has not been shown to bias repair in any way, unlike the I-SceI nuclease. Furthermore, this assay can detect most NHEJ events, including +3 and −3 insertions and deletions. As such, this assay has the sensitivity necessary to be useful for high-throughput screens to identify conditions that change the balance of repair between HR and NHEJ. This could include the functional assessment of mutations of unknown significance in relevant repair proteins such as ATM, ATR, BRCA1/2, and RAD51, among a growing list of proteins known to be involved in repair pathways with variants identified in human cancers. Lastly, we have shown the utility of this assay in identifying potential roles in HR or NHEJ repair for proteins not previously specified in either of the HR or NHEJ repair pathways.

Using this assay, we have identified a distinct difference between the HEK293FT cells and HCT116 cells that could be seen upon transfection of the ssDNA oligo repair template alone. This action resulted in the loss of EGFP signal in HEK293FT cells but not in HCT116 cells. As this loss of signal occurred without the presence of CRISPR/Cas9 and the EGFP-targeting gRNA, it was of note and warranted an explanation. Interestingly, one defining characteristic of HCT116 cells is that they are Mismatch Repair (MMR) deficient. As such, one potential hypothesis to entertain to explain the increase in NHEJ in HEK cells and lack thereof in HCT116 cells might be that the ssDNA oligo induces MMR in some manner (i.e. by introducing mismatches that need to be repaired), which results in the loss of EGFP signal, either mimicking NHEJ or utilizing it in some manner. The data supports this theory in that the addition of both template and gRNA increases the chances of NHEJ further, in an additive manner, in HEK cells but not in HCT116 cells. Furthermore, in HCT116 cells, we see a marked reduction in NHEJ when the template is introduced along with the gRNA. This indicates that without the additive impact of the template on rates of EGFP loss, either by NHEJ or MMR, the proportion of NHEJ observed is reduced as a byproduct of the cells being repaired by HR instead. Additionally, HCT116 cells are markedly devoid of rearrangements, a process that utilizes NHEJ in order to occur. This indicates that MMR deficiency in these cells may have some impact on rates of NHEJ as well. That being said, a secondary potential hypothesis for the increase in NHEJ as a result of the addition of the repair template alone is due to an interferon response within the HEK293FT cells, in reaction to the presence of ssDNA. Such a response can lead to increased DNA damage across the genome, including at the EGFP locus, subsequent repair of which by NHEJ could result in the loss of EGFP signal. However, further study would be needed for validation.

To assess the impact of genetic modulation of repair factors using the RepairSwitch assay we utilized ATM wild-type and knockout cells lines. ATM is a key protein in DSB repair by homologous recombination and is recruited to the site of repair by the MRN complex. Its activity at the site of repair initiates a cascade of phosphorylation that recruits additional repair proteins and initiates signaling pathways that lead to cell cycle arrest so repair can take place. While we saw the expected reduction in rates of HR upon using our assay in the ATM knockout cells, we saw an interesting increase in the rate of NHEJ in these cells as well, as compared to the wild-type. This introduced the possibility that rates of NHEJ may be increased in HR deficient cells to compensate for the loss of effective HR, and warranted further study. When we interrogated this pathway with the use of the ATM inhibitor KU-60019, it further strengthened the hypothesis that this was indeed a compensatory mechanism utilized by the cells that could no longer perform HR. Furthermore, this effect was also observed when DNK-PK, an important component in the NHEJ machinery, was inhibited, showing that the converse is also true. As such, we have shown using the RepairSwitch assay, that both repair pathways, HR and NHEJ, can and will compensate for the loss of the other when necessary.

We next examined pharmacological inhibition of PARP to evaluate its impact on rates of HR and NHEJ. PARP is a well-studied repair factor in the base excision repair (BER) pathway, and does not work in the canonical pathway of either HR or NHEJ. However, recent studies have shown that it may play a role in NHEJ that is as of yet not fully understood (Wang et al. 2006). This is of great interest as it presents potential vulnerabilities that can be exploited by treatment with Olaparib in HR defective patients. These tactics are already being employed in several clinical trials. Our results show a subtle reduction in rates of HR and a correspondingly subtle increase in the rates of NHEJ. Although these data are interesting, and seemingly contradictory to the implication that PARP is involved in NHEJ and not HR, these data do not definitively implicate PARP in having a direct role in either the HR or NHEJ pathways. Repair proteins often have multiple and complex functions that can span a number of pathways. It is possible that PARP has roles in both HR and NHEJ that can explain the results we see and those seen in clinical trials.

The RepairSwitch assay also revealed the relationship between DNA damage repair and the methylation machinery. We have shown that use of a methylated repair template significantly decreases the rates of both HR and NHEJ as compared to an unmethylated repair template in HEK293FT cells but not in HCT116 cells. This indicates the mechanisms by which cells utilize methylated vs unmethylated repair templates is a cell line dependent process.

While it is not clear what role DNMTs have in repair, it has been shown that they are present at the site of repair by IF (Mortusewicz et al. 2005). Our use of Decitabine to inhibit DNMTs results in a dose responsive decrease in the rate of HR and a compensatory increase in the rate of NHEJ in both HEK293FT cells and HCT116 cells. Interestingly, this pattern of loss of HR, and increased utilization of NHEJ for repairing DNA is what is observed in cells with mutations in BRCA1 or BRCA2, or in other HR pathway genes. This suggests that decitabine may induce a “BRCAness” phenotype in which cells treated with decitabine behave functionally as though they have HR deficiency. This would predict that treatment with decitabine would lead to sensitization to PARP inhibitors, which show synthetic lethality in cells with BRCA1 or BRCA2 mutations. Consistent with the prediction, recent studies have shown strong synergy of the DNMT inhibitors with PARP inhibitors for treatment of multiple cancers (Muvarak NE et al, 2016; Pulliam N et al, 2018; Abbotts R et al, 2019; McLaughlin LJ et al, 2020). These observations therefore illustrate the utility of the RepairSwitch assay in identifying highly relevant and meaningful functional alterations in DNA DSB repair.

In conclusion, we have shown the RepairSwitch assay to be an effective tool to better our understanding of the dynamics between homologous recombination and non-homologous end joining. The RepairSwitch assay has much potential to be used as a way to functionally classify mutations in repair proteins, as well as a tool to identify how various therapeutic treatments impact the rates of HR and NHEJ, in the context of both healthy cells and cancer cells. It is the hope that an improved understanding of the DNA repair machinery, in both of these contexts, may lead us to a better understanding of the underlying mechanisms and to new avenues of cancer treatment.

## ACKNOWLEDGMENTS

We thank Jessica Gucwa and the members of the Johns Hopkins Sidney Kimmel Comprehensive Cancer Center Flow Cytometry Core for assistance with the flow cytometry experiments. We thank Dr. Stephen Baylin for kindly providing the DNMT1 and DNMT3B knockout cell lines. We thank Minh Nguyen at Clovis Oncology for kindly providing the CWR22Rv1 ATM knockout cells.

## Notes

### Competing Interest Statement

R.S., W.G.N., and S.Y. are co-inventors in a patent application, managed by the Johns Hopkins Technology Ventures and intellectual property policies, describing the RepairSwitch assay.

## REFERENCES

Abbotts R, Topper MJ, Biondi C, Fontaine D, Goswami R, Stojanovic L, Choi EY, McLaughlin L, Kogan AA, Xia L, Lapidus R, Mahmood J, Baylin SB, Rassool FV. DNA methyltransferase inhibitors induce a BRCAness phenotype that sensitizes NSCLC to PARP inhibitor and ionizing radiation. Proc Natl Acad Sci U S A. 2019 Nov 5;116(45):22609–22618. doi: 10.1073/pnas.1903765116. Epub 2019 Oct 7. PMID: 31591209; PMCID: PMC6842607.

Ahel, Dragana, Zuzana Horejsí, Nicola Wiechens, Sophie E. Polo, Elisa Garcia-Wilson, Ivan Ahel, Helen Flynn, et al. 2009. “Poly(ADP-Ribose)-Dependent Regulation of DNA Repair by the Chromatin Remodeling Enzyme ALC1.” Science 325 (5945): 1240–43. https://doi.org/10.1126/science.1177321.

Ayoub, Nabieh, Anand D. Jeyasekharan, Juan A. Bernal, and Ashok R. Venkitaraman. 2008. “HP1-Beta Mobilization Promotes Chromatin Changes That Initiate the DNA Damage Response.” Nature 453 (7195): 682–86. https://doi.org/10.1038/nature06875.

Baldeyron, Céline, Gaston Soria, Danièle Roche, Adam J. L. Cook, and Geneviève Almouzni. 2011. “HP1alpha Recruitment to DNA Damage by p150CAF-1 Promotes Homologous Recombination Repair.” The Journal of Cell Biology 193 (1): 81–95. https://doi.org/10.1083/jcb.201101030.

Cairns, Bradley R. 2005. “Chromatin Remodeling Complexes: Strength in Diversity, Precision through Specialization.” Current Opinion in Genetics & Development 15 (2): 185–90. https://doi.org/10.1016/j.gde.2005.01.003.

Certo, Michael T., Byoung Y. Ryu, James E. Annis, Mikhail Garibov, Jordan Jarjour, David J. Rawlings, and Andrew M. Scharenberg. 2011. “Tracking Genome Engineering Outcome at Individual DNA Breakpoints.” Nature Methods 8 (8): 671–76. https://doi.org/10.1038/nmeth.1648.

Chou, Danny M., Britt Adamson, Noah E. Dephoure, Xu Tan, Amanda C. Nottke, Kristen E. Hurov, Steven P. Gygi, Monica P. Colaiácovo, and Stephen J. Elledge. 2010. “A Chromatin Localization Screen Reveals Poly (ADP Ribose)-Regulated Recruitment of the Repressive Polycomb and NuRD Complexes to Sites of DNA Damage.” Proceedings of the National Academy of Sciences of the United States of America 107 (43): 18475–80. https://doi.org/10.1073/pnas.1012946107.

Clapier, Cedric R., and Bradley R. Cairns. 2009. “The Biology of Chromatin Remodeling Complexes.” Annual Review of Biochemistry 78: 273–304. https://doi.org/10.1146/annurev.biochem.77.062706.153223.

Cuozzo, Concetta, Antonio Porcellini, Tiziana Angrisano, Annalisa Morano, Bongyong Lee, Alba Di Pardo, Samantha Messina, et al. 2007. “DNA Damage, Homology-Directed Repair, and DNA Methylation.” PLoS Genetics 3 (7): e110. https://doi.org/10.1371/journal.pgen.0030110.

Dawson, Mark A. 2017. “The Cancer Epigenome: Concepts, Challenges, and Therapeutic Opportunities.” Science 355 (6330): 1147–52. https://doi.org/10.1126/science.aam7304.

Glaser, Astrid, Bradley McColl, and Jim Vadolas. 2016. “GFP to BFP Conversion: A Versatile Assay for the Quantification of CRISPR/Cas9-Mediated Genome Editing.” Molecular Therapy. Nucleic Acids 5 (7): e334. https://doi.org/10.1038/mtna.2016.48.

Goldberg, Michal, Manuel Stucki, Jacob Falck, Damien D’Amours, Dinah Rahman, Darryl Pappin, Jiri Bartek, and Stephen P. Jackson. 2003. “MDC1 Is Required for the Intra-S-Phase DNA Damage Checkpoint.” Nature 421 (6926): 952–56. https://doi.org/10.1038/nature01445.

Gonzalo, Susana, Isabel Jaco, Mario F. Fraga, Taiping Chen, En Li, Manel Esteller, and María A. Blasco. 2006. “DNA Methyltransferases Control Telomere Length and Telomere Recombination in Mammalian Cells.” Nature Cell Biology 8 (4): 416–24. https://doi.org/10.1038/ncb1386.

Goodarzi, Aaron A., Angela T. Noon, Dorothee Deckbar, Yael Ziv, Yosef Shiloh, Markus Löbrich, and Penny A. Jeggo. 2008. “ATM Signaling Facilitates Repair of DNA Double-Strand Breaks Associated with Heterochromatin.” Molecular Cell 31 (2): 167–77. https://doi.org/10.1016/j.molcel.2008.05.017.

Grewal, Shiv I. S., and Songtao Jia. 2007. “Heterochromatin Revisited.” Nature Reviews. Genetics 8 (1): 35–46. https://doi.org/10.1038/nrg2008.

Ha, Kyungsoo, Gun Eui Lee, Stela S. Palii, Kevin D. Brown, Yoshihiko Takeda, Kebin Liu, Kapil N. Bhalla, and Keith D. Robertson. 2011. “Rapid and Transient Recruitment of DNMT1 to DNA Double-Strand Breaks Is Mediated by Its Interaction with Multiple Components of the DNA Damage Response Machinery.” Human Molecular Genetics 20 (1): 126–40. https://doi.org/10.1093/hmg/ddq451.

Hickson, Ian, Yan Zhao, Caroline J. Richardson, Sharon J. Green, Niall M. B. Martin, Alisdair I. Orr, Philip M. Reaper, Stephen P. Jackson, Nicola J. Curtin, and Graeme C. M. Smith. 2004. “Identification and Characterization of a Novel and Specific Inhibitor of the Ataxia-Telangiectasia Mutated Kinase ATM.” Cancer Research 64 (24): 9152–59. https://doi.org/10.1158/0008-5472.CAN-04-2727.

Jasin, M. 1996. “Genetic Manipulation of Genomes with Rare-Cutting Endonucleases.” Trends in Genetics: TIG 12 (6): 224–28. https://doi.org/10.1016/0168-9525(96)10019-6.

Kadoch, Cigall, Diana C. Hargreaves, Courtney Hodges, Laura Elias, Lena Ho, Jeff Ranish, and Gerald R. Crabtree. 2013. “Proteomic and Bioinformatic Analysis of Mammalian SWI/SNF Complexes Identifies Extensive Roles in Human Malignancy.” Nature Genetics 45 (6): 592–601. https://doi.org/10.1038/ng.2628.

Larsen, Dorthe Helena, Catherine Poinsignon, Thorkell Gudjonsson, Christoffel Dinant, Mark R. Payne, Flurina J. Hari, Jannie M. Rendtlew Danielsen, et al. 2010. “The Chromatin-Remodeling Factor CHD4 Coordinates Signaling and Repair after DNA Damage.” The Journal of Cell Biology 190 (5): 731–40. https://doi.org/10.1083/jcb.200912135.

Luijsterburg, Martijn S., Christoffel Dinant, Hannes Lans, Jan Stap, Elzbieta Wiernasz, Saskia Lagerwerf, Daniël O. Warmerdam, et al. 2009. “Heterochromatin Protein 1 Is Recruited to Various Types of DNA Damage.” The Journal of Cell Biology 185 (4): 577–86. https://doi.org/10.1083/jcb.200810035.

McLaughlin LJ, Stojanovic L, Kogan AA, Rutherford JL, Choi EY, Yen RC, Xia L, Zou Y, Lapidus RG, Baylin SB, Topper MJ, Rassool FV. Pharmacologic induction of innate immune signaling directly drives homologous recombination deficiency. Proc Natl Acad Sci U S A. 2020 Jul 28;117(30):17785–17795. doi: 10.1073/pnas.2003499117.

Morano, Annalisa, Tiziana Angrisano, Giusi Russo, Rosaria Landi, Antonio Pezone, Silvia Bartollino, Candida Zuchegna, et al. 2014. “Targeted DNA Methylation by Homology-Directed Repair in Mammalian Cells. Transcription Reshapes Methylation on the Repaired Gene.” Nucleic Acids Research 42 (2): 804–21. https://doi.org/10.1093/nar/gkt920.

Mortusewicz, Oliver, Lothar Schermelleh, Joachim Walter, M. Cristina Cardoso, and Heinrich Leonhardt. 2005. “Recruitment of DNA Methyltransferase I to DNA Repair Sites.” Proceedings of the National Academy of Sciences of the United States of America 102 (25): 8905–9. https://doi.org/10.1073/pnas.0501034102.

Muvarak NE, Chowdhury K, Xia L, Robert C, Choi EY, Cai Y, Bellani M, Zou Y, Singh ZN, Duong VH, Rutherford T, Nagaria P, Bentzen SM, Seidman MM, Baer MR, Lapidus RG, Baylin SB, Rassool FV. Enhancing the Cytotoxic Effects of PARP Inhibitors with DNA Demethylating Agents - A Potential Therapy for Cancer. Cancer Cell. 2016 Oct 10;30(4):637–650. doi: 10.1016/j.ccell.2016.09.002.

O’Hagan, Heather M. 2014. “Chromatin Modifications during Repair of Environmental Exposure-Induced DNA Damage: A Potential Mechanism for Stable Epigenetic Alterations.” Environmental and Molecular Mutagenesis 55 (3): 278–91. https://doi.org/10.1002/em.21830.

Okano, M., D. W. Bell, D. A. Haber, and E. Li. 1999. “DNA Methyltransferases Dnmt3a and Dnmt3b Are Essential for de Novo Methylation and Mammalian Development.” Cell 99 (3): 247– 57. https://doi.org/10.1016/s0092-8674(00)81656-6.

Peng, Jamy C., and Gary H. Karpen. 2008. “Epigenetic Regulation of Heterochromatic DNA Stability.” Current Opinion in Genetics & Development 18 (2): 204–11. https://doi.org/10.1016/j.gde.2008.01.021.

Pierce, A. J., R. D. Johnson, L. H. Thompson, and M. Jasin. 1999. “XRCC3 Promotes Homology-Directed Repair of DNA Damage in Mammalian Cells.” Genes & Development 13 (20): 2633–38. https://doi.org/10.1101/gad.13.20.2633.

Pierce, Andrew J., and Maria Jasin. 2005. “Measuring Recombination Proficiency in Mouse Embryonic Stem Cells.” In Molecular Toxicology Protocols, edited by Phouthone Keohavong and Stephen G. Grant, 373–84. Totowa, NJ: Humana Press. https://doi.org/10.1385/1-59259-840-4:373.

Polo, Sophie E., Abderrahmane Kaidi, Linda Baskcomb, Yaron Galanty, and Stephen P. Jackson. 2010. “Regulation of DNA-Damage Responses and Cell-Cycle Progression by the Chromatin Remodelling Factor CHD4.” The EMBO Journal 29 (18): 3130–39. https://doi.org/10.1038/emboj.2010.188.

Price, Brendan D., and Alan D. D’Andrea. 2013. “Chromatin Remodeling at DNA Double-Strand Breaks.” Cell 152 (6): 1344–54. https://doi.org/10.1016/j.cell.2013.02.011.

Pulliam N, Fang F, Ozes AR, Tang J, Adewuyi A, Keer H, Lyons J, Baylin SB, Matei D, Nakshatri H, Rassool FV, Miller KD, Nephew KP. An Effective Epigenetic-PARP Inhibitor Combination Therapy for Breast and Ovarian Cancers Independent of BRCA Mutations. Clin Cancer Res. 2018 Jul 1;24(13):3163–3175. doi: 10.1158/1078-0432.CCR-18-0204.

Russo, Giusi, Rosaria Landi, Antonio Pezone, Annalisa Morano, Candida Zuchegna, Antonella Romano, Mark T. Muller, Max E. Gottesman, Antonio Porcellini, and Enrico V. Avvedimento. 2016. “DNA Damage and Repair Modify DNA Methylation and Chromatin Domain of the Targeted Locus: Mechanism of Allele Methylation Polymorphism.” Scientific Reports 6 (September): 33222. https://doi.org/10.1038/srep33222.

Seluanov, Andrei, Zhiyong Mao, and Vera Gorbunova. 2010. “Analysis of DNA Double-Strand Break (DSB) Repair in Mammalian Cells.” Journal of Visualized Experiments: JoVE, no. 43 (September). https://doi.org/10.3791/2002.

Sishc, Brock J., and Anthony J. Davis. 2017. “The Role of the Core Non-Homologous End Joining Factors in Carcinogenesis and Cancer.” Cancers 9 (7). https://doi.org/10.3390/cancers9070081.

Smeenk, Godelieve, Wouter W. Wiegant, Hans Vrolijk, Aldo P. Solari, Albert Pastink, and Haico van Attikum. 2010. “The NuRD Chromatin-Remodeling Complex Regulates Signaling and Repair of DNA Damage.” The Journal of Cell Biology 190 (5): 741–49. https://doi.org/10.1083/jcb.201001048.

Smerdon, M. J. 1991. “DNA Repair and the Role of Chromatin Structure.” Current Opinion in Cell Biology 3 (3): 422–28. https://doi.org/10.1016/0955-0674(91)90069-b.

Smerdon, M. J., T. D. Tlsty, and M. W. Lieberman. 1978. “Distribution of Ultraviolet-Induced DNA Repair Synthesis in Nuclease Sensitive and Resistant Regions of Human Chromatin.” Biochemistry 17 (12): 2377–86. https://doi.org/10.1021/bi00605a020.

Soria, Gaston, Sophie E. Polo, and Geneviève Almouzni. 2012. “Prime, Repair, Restore: The Active Role of Chromatin in the DNA Damage Response.” Molecular Cell 46 (6): 722–34. https://doi.org/10.1016/j.molcel.2012.06.002.

Stewart, Grant S., Bin Wang, Colin R. Bignell, A. Malcolm R. Taylor, and Stephen J. Elledge. 2003. “MDC1 Is a Mediator of the Mammalian DNA Damage Checkpoint.” Nature 421 (6926): 961–66. https://doi.org/10.1038/nature01446.

Stresemann, Carlo, and Frank Lyko. 2008. “Modes of Action of the DNA Methyltransferase Inhibitors Azacytidine and Decitabine.” International Journal of Cancer. Journal International Du Cancer 123 (1): 8–13. https://doi.org/10.1002/ijc.23607.

Stucki, Manuel, Julie A. Clapperton, Duaa Mohammad, Michael B. Yaffe, Stephen J. Smerdon, and Stephen P. Jackson. 2005. “MDC1 Directly Binds Phosphorylated Histone H2AX to Regulate Cellular Responses to DNA Double-Strand Breaks.” Cell 123 (7): 1213–26. https://doi.org/10.1016/j.cell.2005.09.038.

Sun, Yingli, Xiaofeng Jiang, Ye Xu, Marina K. Ayrapetov, Lisa A. Moreau, Johnathan R. Whetstine, and Brendan D. Price. 2009. “Histone H3 Methylation Links DNA Damage Detection to Activation of the Tumour Suppressor Tip60.” Nature Cell Biology 11 (11): 1376–82. https://doi.org/10.1038/ncb1982.

Unterberger, Alexander, Stephen D. Andrews, Ian C. G. Weaver, and Moshe Szyf. 2006. “DNA Methyltransferase 1 Knockdown Activates a Replication Stress Checkpoint.” Molecular and Cellular Biology 26 (20): 7575–86. https://doi.org/10.1128/MCB.01887-05.

Vriend, Lianne E. M., Maria Jasin, and Przemek M. Krawczyk. 2014. “Assaying Break and Nick-Induced Homologous Recombination in Mammalian Cells Using the DR-GFP Reporter and Cas9 Nucleases.” Methods in Enzymology 546: 175–91. https://doi.org/10.1016/B978-0-12-801185-0.00009-X.

Wang, Minli, Weizhong Wu, Wenqi Wu, Bustanur Rosidi, Lihua Zhang, Huichen Wang, and George Iliakis. 2006. “PARP-1 and Ku Compete for Repair of DNA Double Strand Breaks by Distinct NHEJ Pathways.” Nucleic Acids Research 34 (21): 6170–82. https://doi.org/10.1093/nar/gkl840.

Xu, G. L., T. H. Bestor, D. Bourc’his, C. L. Hsieh, N. Tommerup, M. Bugge, M. Hulten, X. Qu, J. J. Russo, and E. Viegas-Péquignot. 1999. “Chromosome Instability and Immunodeficiency Syndrome Caused by Mutations in a DNA Methyltransferase Gene.” Nature 402 (6758): 187– 91. https://doi.org/10.1038/46052.

Ziv, Yael, Dana Bielopolski, Yaron Galanty, Claudia Lukas, Yoichi Taya, David C. Schultz, Jiri Lukas, Simon Bekker-Jensen, Jiri Bartek, and Yosef Shiloh. 2006. “Chromatin Relaxation in Response to DNA Double-Strand Breaks Is Modulated by a Novel ATM- and KAP-1 Dependent Pathway.” Nature Cell Biology 8 (8): 870–76. https://doi.org/10.1038/ncb1446.

