## Supplementary Figures for "RepairSwitch: simultaneous functional assessment of homologous recombination vs end joining DNA repair pathways in living cells"

SUPPLEMENTARY FIGURE 1.

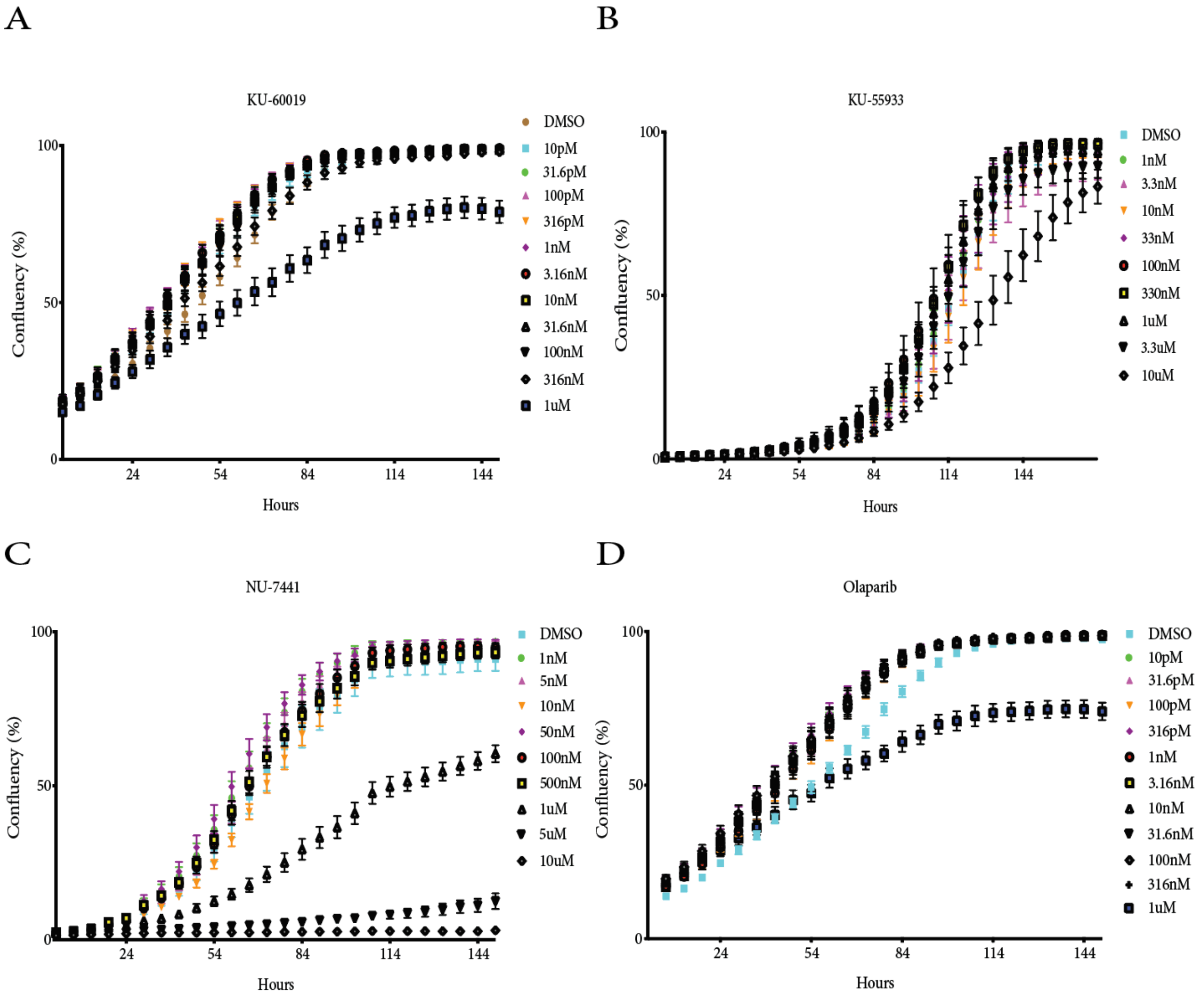


**Figure S1. Incucyte Proliferation Assay.** Growth curves for (A) KU-60019 ATM inhibitor (B) KU-55933 ATM inhibitor (C) NU-7441 DNA-PK inhibitor and (D) Olaparib PARP inhibitor, to ascertain toxicity and growth inhibition.

SUPPLEMENTARY FIGURE 2.

A

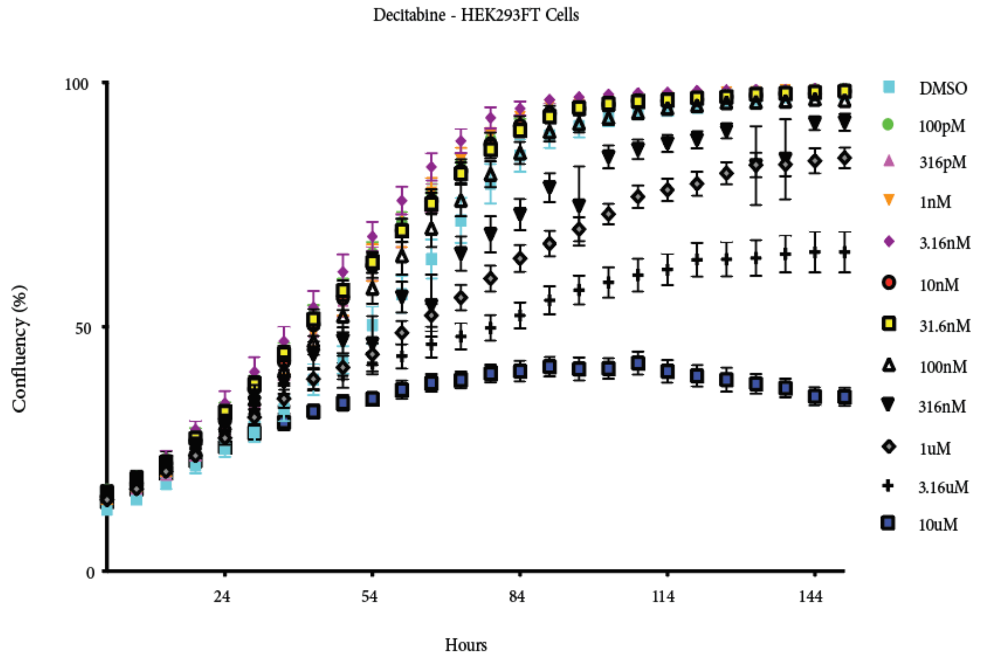


B

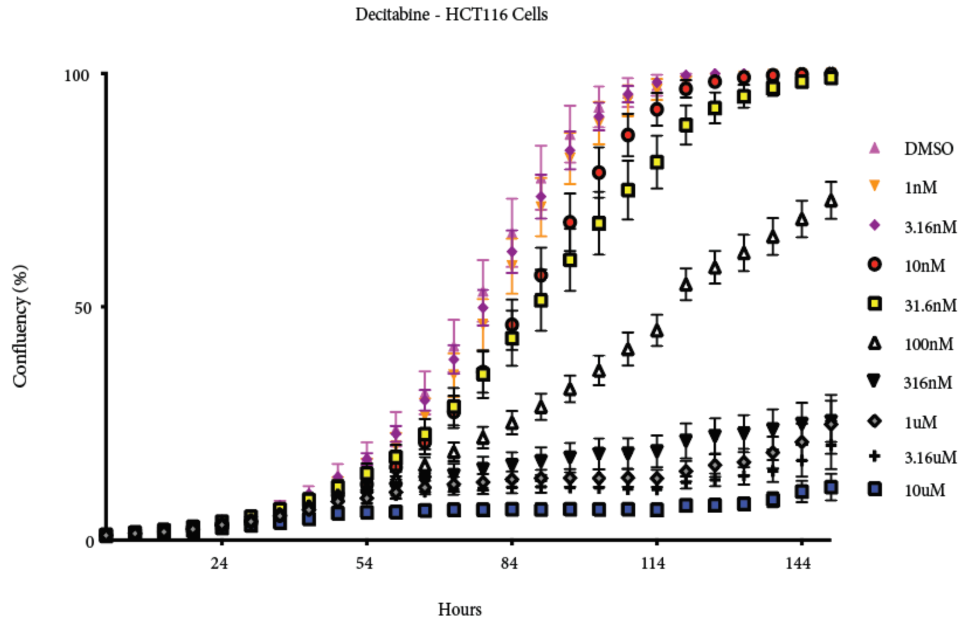


**Figure S2. Incucyte Proliferation Assay.** Incucyte was performed on both (A) HEK293FT cells and (B) HCT116 cells to ascertain the toxicity and growth inhibition post Decitabine treatment.
